# Measuring vision using innate behaviours in mice with intact and impaired retina function

**DOI:** 10.1101/520247

**Authors:** R. Storchi, J. Rodgers, M. Gracey, F.P. Martial, J. Wynne, S. Ryan, C.J. Twining, T.F. Cootes, R. Killick, R.J. Lucas

## Abstract

Measuring vision in rodents is a critical step for understanding vision, improving models of human disease, and developing therapies. Established behavioural tests for perceptual vision, such as the visual water task, rely on learning. The learning process, while effective for sighted animals, can be laborious and stressful in animals with impaired vision, requiring long periods of training. Current tests that that do not require training are based on sub-conscious, reflex responses (e.g. optokinetic nystagmus) that don’t require involvement of visual cortex and higher order thalamic nuclei. A potential alternative for measuring vision relies on using visually guided innate defensive responses, such as escape or freeze, that involve cortical and thalamic circuits. In this study we address this possibility in mice with intact and degenerate retinas. We first develop automatic methods to detect behavioural responses based on high dimensional tracking and changepoint detection of behavioural time series. Using those methods, we show that visually guided innate responses can be elicited using parametisable stimuli, and applied to describing the limits of visual acuity in healthy animals and discriminating degrees of visual dysfunction in mouse models of retinal degeneration.

## Introduction

Rodents, and in particular mice, are increasingly applied to understanding the physiology and neural computations underlying vision. Rodent models of ocular diseases are also an important tool to understand disease and trial therapies. A great body of work has been dedicated to developing rodent models that capture the critical aspects of human diseases and there are currently >100 different types of visually impaired mouse (Chang et al., 2005). A particular focus has been devoted to using these models to develop treatments based on optogenetics, stem cell therapies and gene therapies for such incurable conditions as retinitis pigmentosa and Leber congenital amaurosis (Auricchio et al., 2017; Busskamp et al., 2012; Takahashi et al., 2018).

A crucial step in describing degeneration and establishing therapeutic efficacy in such pre-clinical studies is assessment of visual function, commonly by electrophysiological and/or behavioural approaches. Electrical recordings of neural activity in the retina and in the brain can be very useful in revealing physiological responses to visual stimuli. However, they don't directly measure integrated visual performance or perceptual components of vision. Many commonly used behavioural assays of visual function, such as pupil constriction (Hattar et al., 2003), optokinetic and optomotor reflexes (Abdejalil et al., 2005; Stahl, 2004), suffer from the same weakness; being sub-conscious, reflex responses whose activity is only indirectly related to perceptual vision. All current tests that allow measurement of perceptual, cortically modulated vision rely on learning (Busse et al., 2011; Morton et al., 2006; Prusky and Douglas, 2004). The most commonly used, the visual water task, has been effectively and widely employed to measure spatial acuity and brightness discrimination in mice with intact visual systems. However in animal models of moderate or severe retinal degeneration this task can only be painstakingly learnt through several weeks of intensive training (Thyagarajan et al., 2010). This process is time consuming and can be significantly stressful for the animals (Vorhees and Williams, 2014). Furthermore it can introduce additional sources of variability including the effects of repeated handling (Hurst and West, 2010) and the learning process itself (Coppens et al., 2010; Flagel et al., 2009).

The aim of this work is to develop a rapid assay of visual stimulus detection, suitable for mice with intact and degenerate retinas, which doesn’t rely on training. To meet these requirements we investigate the possibility of using innate behaviours instead of learned tasks. Visually evoked changes in exploratory behaviour have been applied to this problem (Cehajic-Kapetanovic et al., 2015; De Silva et al., 2017), but such tests are relatively low throughput and we were interested in the possibility of exploiting recent advances in understanding of rapid defensive behaviours as an alternative. Behaviourally salient visual stimuli, such as a looming dark spot (De Franceschi et al., 2016; Yilmaz and Meister, 2013) or a full field flash of light (Liang et al., 2015), can reliably trigger different behavioural responses. Several studies aimed at elucidating the circuits involved in controlling these behaviours revealed a substantial involvement of primary visual cortex (Liang et al., 2015; Zhao et al., 2014) and first and higher order thalamic nuclei (Shang et al., 2015b; Wei et al., 2015b) as well as spatial memory (Vale et al., 2017). This indicates that these behaviours are not simply sub-conscious reflexes and instead rely on processing of higher order image forming pathways and integration with the limbic system. They are rapidly induced and short lived, meaning that they can, in theory, be evoked multiple times in a single experimental session. However so far such innate responses haven’t been used systematically to assay visual ability, nor have they been tested in animals with mild or severe visual impairments. Therefore it is not clear to what extent these responses could be used to discriminate different degrees of retinal function.

We show that relating induction of innate responses to features of visual stimuli represents an efficient strategy for measuring the limits of visual acuity in mice with an intact visual system. Importantly innate responses are partially maintained in animals with impaired vision and can be used to discriminate different levels of vision loss.

## Results

### Behavioural Responses in Visually Intact Mice

We first set out to apply innate responses to measure visual capabilities in mice with an intact visual system. To this end, we developed a simple algorithm for tracking many landmarks on an animal’s body in order to improve our ability to detect and characterise behavioural responses in an open field arena (**Figure 1a,b**, see Methods for details). For each pair of consecutive frames the change in location of landmarks was used to calculate speed of motion (**Figure 1b**, lower panels), and the distribution of velocities across landmarks divided into quantiles (**Figure 1b**, upper panels) and used to generate a multidimensional time series representing the movements of different body parts (**Figure 1c**). By plotting distinct behavioural time series for each quantile we could then detect occasions in which movement was restricted to parts of the body (e.g. head movements or rearing; **Figure 1b**, middle panel), as well as full body movements (e.g. locomotion; **Figure 1b**, right panel).

**Figure 1:**
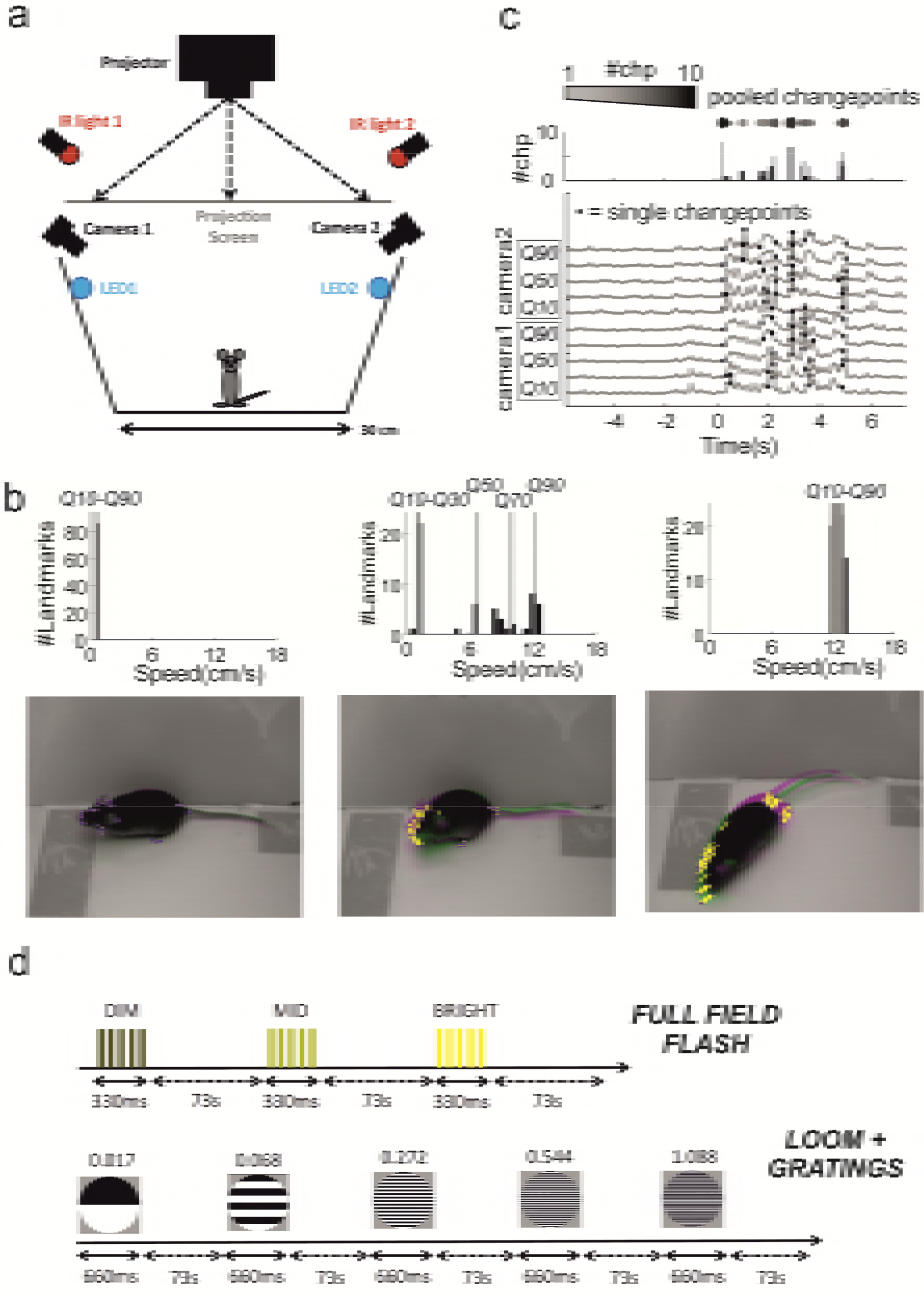
Behavioural arena, data analyses and experimental protocols. **a**) Schematic of the open field arena used for all tests (dimension: 30×30cm). Power LEDs were used to provide diffuse illumination for the flashes while a rear projection screen was used to deliver the looming stimuli. **b**) Images of a mouse in the arena (bottom) over successive frames (superimposed pink and green images) during periods of quiescence (left), head movement (middle) and locomotion (right). Landmarks were automatically identified on the mouse’s body (blue dots on images at bottom) and the speed of each landmark calculated as proportional to the change in position from previous frame (visualised as yellow lines). Distribution of speed across landmarks was then calculated for each frame (top panels) and 10^th^, 30^th^, 50^th^, 70^th^, 90^th^ speed quantiles (respectively Q10 to Q90) used for subsequent analyses. **c**) Behavioural time series showing velocity of landmarks (Q10-90) extracted from two cameras (camera 1 and 2, bottom panel; visual stimulus at time 0). Changepoint detection was run independently on each series (single changepoints depicted by black dots) and identified changepoints were then summed for each frame across those series (top panels, “pooled changepoints”) and used for changepoint statistics throughout the study. **d**) Schematic of the two stimulation protocols used for full field flash and pattern detection. In both protocols the order of each stimulus is block randomised (note: each block is represented by the list of all the distinct stimuli for that protocol).

We considered two ways of using the tracking data to measure behavioural responses to a visual stimulus. In the simplest, we could use the estimates of landmark speed to identify increases or decreases in movement. In the second, we established changepoint analysis as a method to identify responses, and to quantify the frequency and reliability of visually induced behaviours. Changepoint analysis is used to detect if and when the properties of a time series change. By applying this technique to the behavioural time series, we can identify when a mouse’s behaviour changes. We reasoned that if visual stimuli affect behaviour then we would expect higher rates of behavioural changes to occur after the onset of those stimuli (shown in **Figure 1c**). Thus we used Pruned Exact Linear Time algorithm (PELT)(Killick et al., 2012) to detect changes in speed in our behavioural time series. This method is computationally efficient and can analyse dozens of our time series in less than a second. The PELT algorithm uses a penalty term to prevent overfitting changepoints (see Methods). This penalty term determines how much a changepoint must improve the model fit in order to be included. Small values result in a high rate of changepoints and are suitable to detect small changes in movements while larger values will only detect more substantial changes. In order to identify a suitable value for our data we investigated how the penalty value affects changepoint detection using the CROPS algorithm (Haynes et al., 2017) and knee estimation (Muggeo, 2003) (see Methods for details and **Supplementary Figure 1b,c**).

We designed two sets of stimuli aimed at probing different levels of visual capabilities. For the first we presented the animals with sudden changes in ambient light (bursts of 5×30ms flashes over 330ms, **Figure 1d**, “full field flash”) at three different magnitudes (see **Table 1** for calibrated background and flash intensities). Detection of this stimulus relies on full field contrast sensitivity and does not require fine spatial discrimination. For the second set we generated a modified looming stimulus where the dark enlarging spot was replaced by a static grating whose average intensity was isoluminant with the grey background (**Table 2**; Spatial Frequency = 0.017, 0.068, 0.272, 0.544, 1.088 cycles/degree; stimulus duration: 660ms; looming speed = 66deg/s; Inter-Stimulus-Interval = 73 seconds; **Figure 1d**, “loom+gratings”). By changing the spatial frequency of the grating we could then use this stimulus to probe spatial acuity. As background illumination in both experiments we used low photopic light (see **Table 1&2**).

**Table 1:** Background irradiance (log_10_ photon/cm^2^/s).

|  | S-cone opsin | Melanopsin | Rhodopsin | M-cone opsin |
| --- | --- | --- | --- | --- |
| **Full Field Flash Contrast Response** | 10.0552 | 12.765 | 12.8345 | 13.0188 |
| **Gratings Looming for Spatial Acuity** | 10.6106 | 13.2171 | 13.2868 | 13.4712 |
| **Full Field Flash Contrast Response (RD1 mice)** | 9.16516 | 12.0543 | 12.1219 | 12.2971 |

**Table 2:** Michelson Contrast (%).

| **Full Field Contrast Response** | S-cone opsin | Melanopsin | Rhodopsin | M-cone opsin |
| --- | --- | --- | --- | --- |
| DIM | 32.8793 | 44.1403 | 35.2123 | 11.2034 |
| MID | 86.1193 | 89.1086 | 84.8028 | 56.2394 |
| BRIGHT | 99.4891 | 99.5334 | 99.3131 | 97.0982 |
| **Gratings Looming for Spatial Acuity\*** | S-cone opsin | Melanopsin | Rhodopsin | M-cone opsin |
| White vs Grey | 35.254 | 35.254 | 35.254 | 35.254 |
| Black vs Grey | -87.113 | -87.113 | -87.113 | -87.113 |
| White vs Black | 93.617 | 93.617 | 93.617 | 93.617 |
| **Full Field Flash Contrast Response (RD1 mice)** | S-cone opsin | Melanopsin | Rhodopsin | M-cone opsin |
| DIM | 79.1772 | 80.2339 | 73.7149 | 39.9271 |
| MID | 97.9658 | 97.6759 | 96.6435 | 87.1301 |
| BRIGHT | 99.9339 | 99.9088 | 99.8661 | 99.4359 |

\*Michelson Contrast measured between white area of the gratings and grey background (White vs Grey), black area of the gratings and the grey background (Black vs Grey), white and black area of the gratings (White vs Black).

The mice expressed a variety of responses that differed both quantitatively and qualitatively. The full field stimulus evoked partial body movements, typically head movements or rearing, or full body changes in locomotor activity (respectively **movie1&2**). The looming stimulus instead reduced the animal activity both when engaged in full body movements such as during locomotion and when the animal was performing more stationary exploration (respectively **movie3&4**). Therefore the responses to flashes and looming were qualitatively different and consisted respectively of an increase and a decrease in movement speed.

The response to full field stimuli was apparent in analyses based upon speed of movement and changepoint detection. Thus, velocity of movement across landmarks increased following presentation of mid and bright, but not dim, flashes (**Figure 2a**; p = 1, 0.006, 4*10^−7^ respectively dim, mid, bright flash; n = 84 trials from 12 animals, signtest). The average responses to highest intensity were substantially larger than to mid intensity levels (p = 0.013, z = 2.48, n = 84 “mid” & “bri” trials, ranksum test). We also found an increase in the number of changepoints after the stimulus onset compared to baseline activity, and this effect was flash intensity dependent (**Figure 2b**, p < 0.0001, df = 2, χ^2^ = 27.57, n = 84, kruskalwallis test). We wondered whether the larger average effects associated with high intensity was also due to an increase in reliability leading to more effective summation across trials. In order to measure reliability of these responses we compared the rate of changepoints before and after stimulus onset on a single trial basis (calculated over time windows of 0.6s duration). The rate increased in 74% trials and decreased in 11% trials for the highest intensity while for the mid-level intensity we found increments and decrements respectively in 45% and 27% trials (**Figure 2c**).

**Figure 2:**
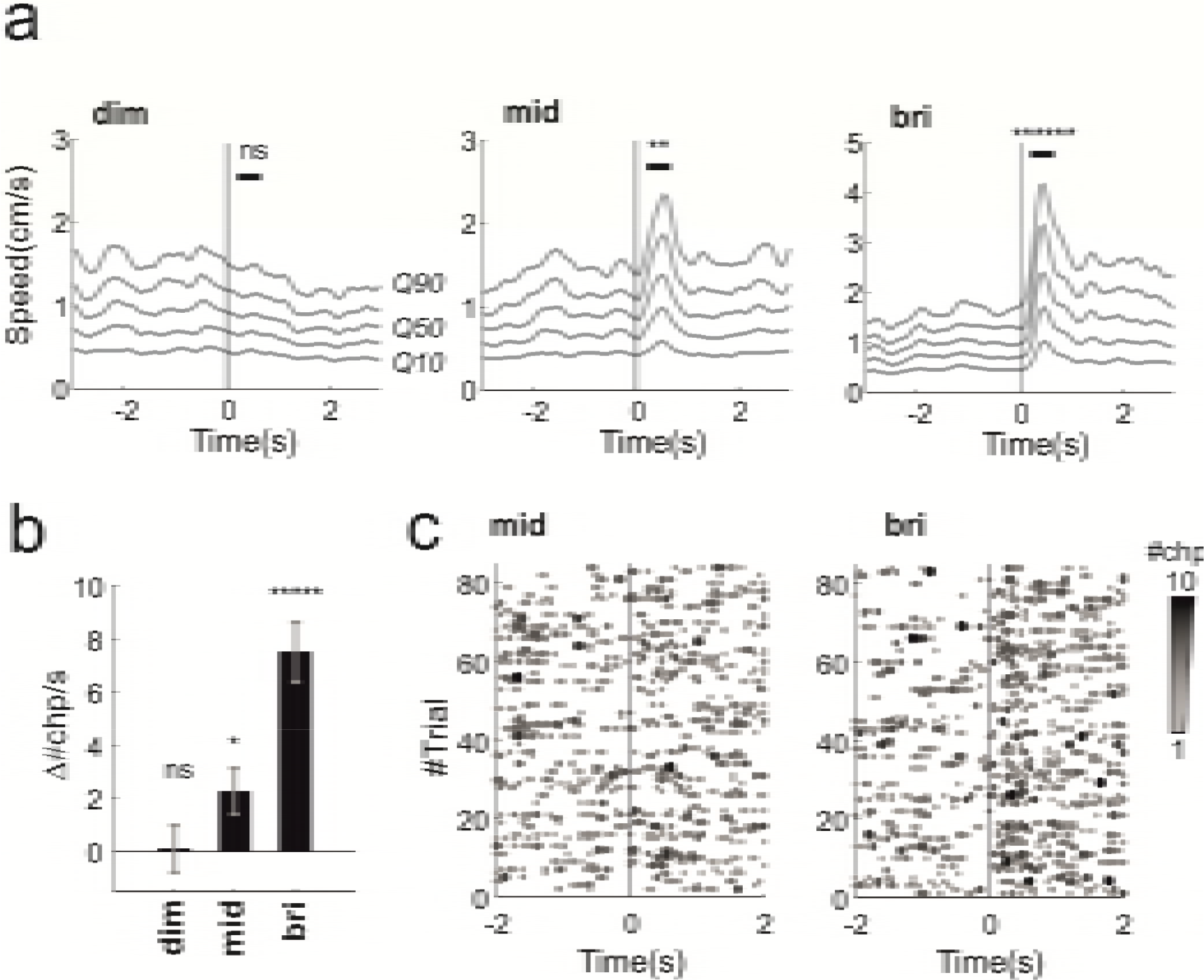
Behavioural responses to full field flashes in visually intact animals. **a**) Movement responses to dim, mid and bright flashes (respectively left, middle and right panels; visual stimulus at time 0). Each grey line represents a different quantile of the speed distribution (10^th^, 30^th^, 50^th^, 70^th^, 90^th^ quantiles, respectively Q10 to Q90, averaged across the two cameras used to track behaviour, see also **Figure 1a-c** and Methods). **b**) Difference between changepoint rate after and before stimulus onset (calculated as *Δ#chp/s = #chp/s (pre) - #chp/s (post)*, see Methods; the rates were calculated in time windows of 0.6s). **c**) Behavioural changepoints for individual trials (stacked along the y-axis) for mid and bright flashes (84 trials collected from 12 animals; each animals recorded for 7 trials; visual stimulus at time 0; ranksum test). For a given trial the number of changepoints occurring at the same time frame are colour coded gray-to-black as exemplified in **Figure 1c.** (“pooled changepoints”). * p < 0.05, ** p < 0..01, *** p < 0.005, **** p < 0.001, ***** p < 0.0005, ****** p < 0.0001, ns = not significant.

The looming stimuli elicited freezing-like reductions in activity across a wide range of frequencies up to 0.544 cpd but not at 1.088 cycles/degrees (p = 0.021, 5*10^−5^, 5*10^−5^, 0.0063, 0.558 respectively 0.017, 0.068, 0.272, 0.544, 1.088 cycles/degrees; n = 91,91,91,105,105 trials from 15 animals, signtest; **Figure 3a**). The changepoint analysis recapitulated those results (**Figure3b**). The most reliable response occurred with the intermediate spatial frequency gratings 0.068 and 0.272 cycles/degree (**Figure 3c**) in which changepoint rate increased respectively in 61% and 65% trials and decreased in 30% and 26%. The highest reproducibility at 0.272 cycles/degree is consistent with peak in contrast sensitivity recorded with visual water task (Prusky and Douglas, 2004). Importantly, the threshold acuity found here (0.544 cpd) is higher than that defined in cortically lesioned animals (<0.3 cycles/degree with visual water task (Douglas et al., 2005)) indicating that visual cortex is involved in driving these behavioural responses. This possibility is also consistent with the described positive cortical modulation of looming evoked responses in superior colliculus (Zhao et al., 2014).

**Figure 3:**
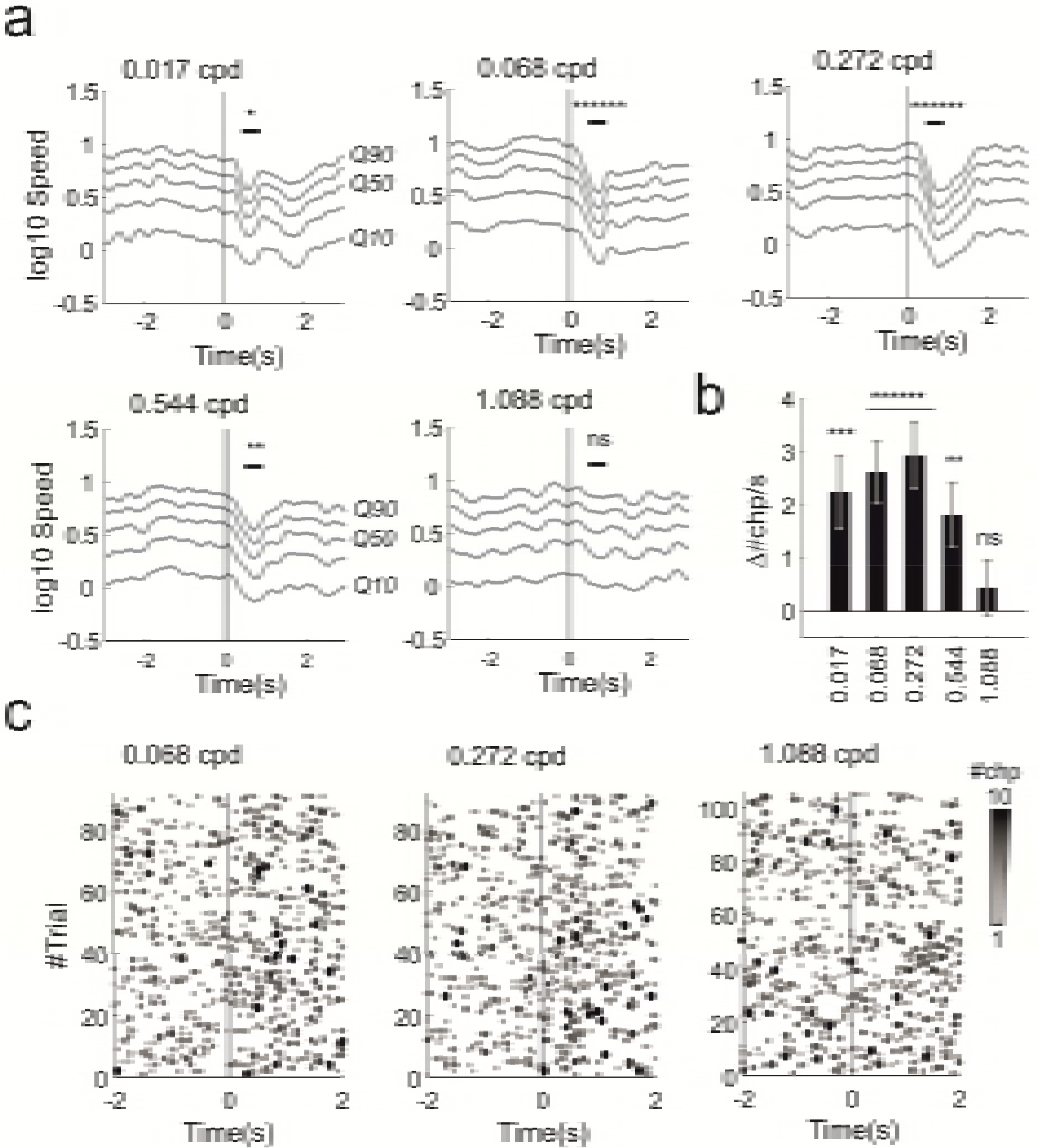
Behavioural responses to looming+gratings in visually intact animals. **a**) Movement responses (in log scale for 10^th^, 30^th^, 50^th^, 70^th^, 90^th^ quantiles - respectively Q10 to Q90 - averaged across two cameras) to all spatial frequencies tested for the spatial acuity stimulus represented in **Figure 1d** (the spatial frequency for each panel is indicated in cycles/degree in the top left corner of each panel; visual stimulus at time 0). Reduction in movement was significant for all frequencies apart from the highest (1.088 cycles/degree). **b**) Differences in changepoint rate before and after stimulus onset across all spatial frequencies tested (mean±sem; spatial frequencies are indicated as cycles/dregrees on the x-axis). **c**) Behavioural changepoints for individual trials at different spatial frequencies (0.068, 0.272, 1.088 cycles/degree; 91, 91, 105 trials, 7 trials/animal; visual stimulus at time 0). * p < 0.05, ** p < 0..01, *** p < 0.005, **** p < 0.001, ***** p < 0.0005, ****** p < 0.0001, ns = not significant.

These results indicate that we can effectively use parametrised stimuli to measure different visual abilities, ranging from coarse light detection to noise-limited spatial acuity, in mice with intact retinas. Interestingly reactions to full field flashes are of opposite sign compared with looming gratings, providing a novel case in which vision guiding of spontaneous behaviour is diverse yet systematic.

### Behavioural Responses in Mouse Models of Retinal Degeneration

The use of innate responses as a tool to measure different levels of visual function in mice with intact visual system suggest that these responses could also be used to probe residual visual function in mouse models of retinal degeneration.

To investigate this possibility we first repeated our behavioural tests in rd1 mice where a mutation of the Pde6b gene substantially disrupts the rod phototransduction cascade (Lavail and Sidman, 1974). These animals represent a model of severe retinal degeneration characterised by fast onset, in which the rod phototransduction cascade is non-functional from birth and cones undergo rapid progressive degeneration (Chang et al., 2002). Adult rd1 mice (3-6 months) undergoing intensive training with visual water task (~3 weeks) can still learn a coarse dark/bright discrimination (Thyagarajan et al., 2010), which is at least partly reliant on inner retina photoreception (Brown et al., 2012), while detection of spatially structured patterns is abolished (Thyagarajan et al., 2010). Previous work has shown that rd1 animals retain some innate light aversion however this behaviour is expressed only after prolonged light exposure (~10 minutes; (Semo et al., 2010)). Moreover, open field spontaneous exploratory behaviour is unaffected by exposure to bright light (Cehajic-Kapetanovic et al., 2015; Lagali et al., 2008). It is currently unknown whether their residual visual functionality can drive more transient behavioural responses resembling those we observed in visually intact animals; and, if that were the case, whether it is possible, based on these responses, to track the progress of visual degeneration.

To answer these questions we selected two cohorts of adult rd1 mice associated with different age groups hereinafter defined as “young” and “old” rd1 (n = 8,12 animals; 10.2±0sd and 24±1.75sd weeks). It was previously shown that cone density undergoes substantial changes between these ages (Lin et al., 2009). The gradual loss of photoreceptor inputs then results in plastic changes and aberrant rhythmic activity at the level of the retinal ganglion cells (Zeck, 2016) targeting both visual thalamus and superior colliculus. Therefore behavioural responses, if at all present, would be expected to be more detectable in young animals.

As a first test we used the same full field flashes delivered to control mice but we reduced the background light intensity in order to maximise effectiveness of the stimulus (see **Table 1&2** for calibrated intensities). The highest intensity flash evoked a transient increase in activity in both age groups (**movie5&6**, **Figure 4a**; p = 0.011 and 0.0375, n = 56 and 84 trials, signtest). The responses could be driven by residual cones, that consistently with previous reports (Lin et al., 2009) we found also in the “old” group (**Figure 4b**) or by intrinsically photosensitive retinal ganglion cells, as melanopsin signalling is largely preserved in these animals (Procyk et al., 2015; Zhang et al., 2008), either by direct input to visual targets in the brain or by modulation of cone signalling (Milosavljevic et al., 2018). Lower intensities did not evoke significant responses (“young”: p = 0.894 and 0.504, n = 56 trials; “old”: p = 0.230 and 0.063, n = 84 trials; signtest for dim and mid intensity).

**Figure 4:**
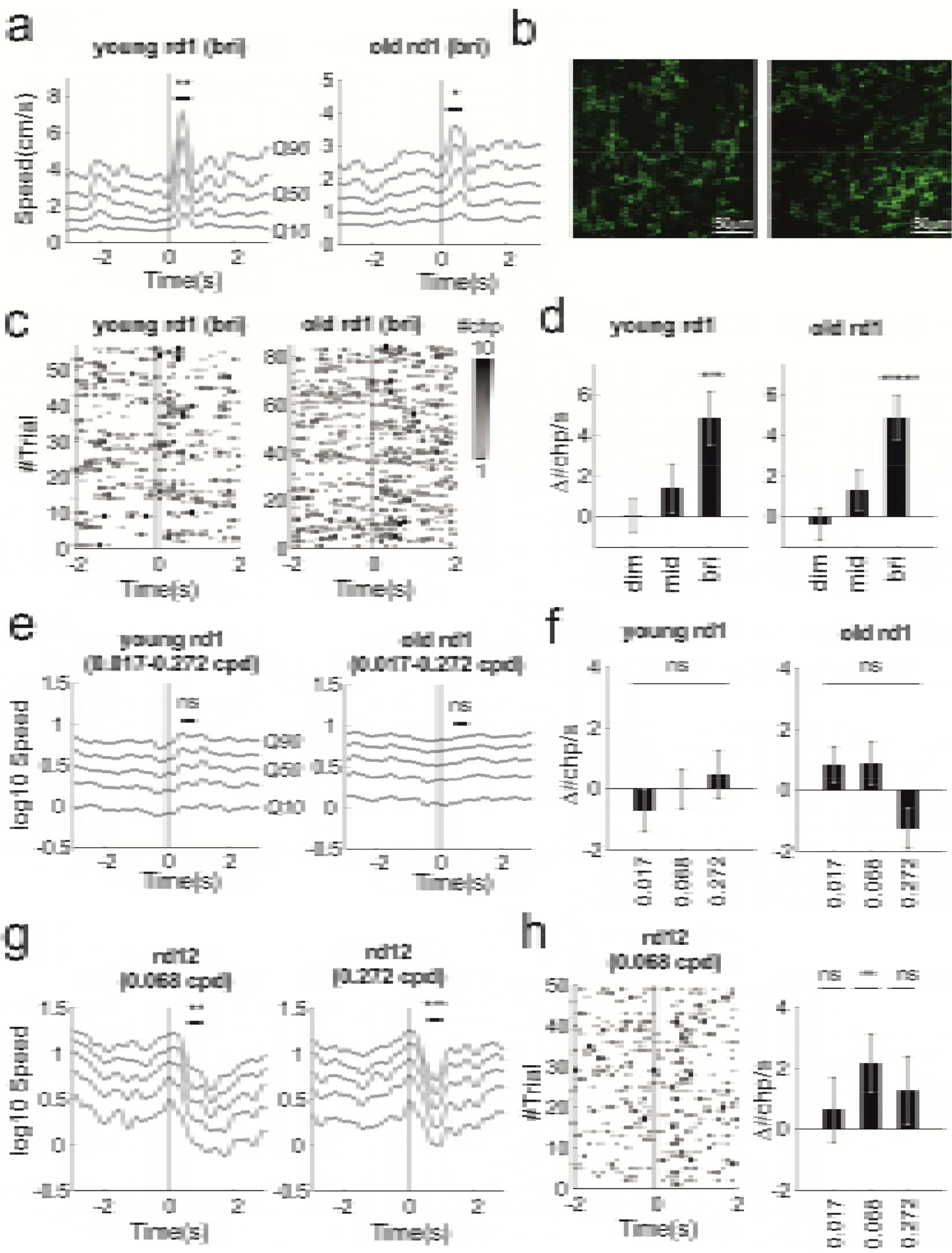
Behavioural responses to full field flashes and looming+gratings in mouse models of retinal degeneration. **a**) Increase in speed after bright flash for young and old rd1 animals (respectively left and right panels). **b**) Immunostaining for s-cone opsin in “old” rd1 retinas (n = 2 retinas, respectively left and right panel). Examination of whole mount retinas by two photon microscopy reveals presence of s-cone opsin in far ventral regions. **c**) Behavioural changepoint for individual trials under bright flash stimuli for “young” and “old” animals (respectively left and right panels; 56, 84 trials collected from 8, 12 animals; each animals recorded for 7 trials; visual stimulus at time 0). **d**) Increase in changepoint rate (mean±sem) as function of flash intensity. **e**) Looming did not evoke changes in speed in both groups. Speed, represented in log scale, was averaged across the spatial frequencies tested (0.017, 0.068 and 0.272 cycles/degree). **f**) Changepoint rates did not significantly change after stimulus onset at all spatial frequencies tested (0.017, 0.068, 0.272 cycles/deg; mean±sem). **g**) rd12 expressed significant reductions in movement at 0.068 cycles/deg (left panel) and at 0.272 cycles/deg (right panel). **h**) Behavioural changepoints for individual trials during looming gratings at 0.068 cycles/deg reveals repeatable responses (left). Changepoint rates as function of spatial frequencies (right panel; mean±sem). * p < 0.05, ** p < 0..01, *** p < 0.005, **** p < 0.001, ***** p < 0.0005, ****** p < 0.0001, ns = not significant.

The response to the highest intensity flash was significantly larger in the young cohort indicating that this simple test allows to detect differences in coarse light detection among the two groups (p = 0.022, z = 2.3, n = 56 and 84, respectively “young” and “old” trials, ranksum test). Compared with control animals trial-to-trial reproducibility was reduced in both groups as changepoint rate increased in 57% (52%) and decreased in 16% (23%) of the trials (respectively young and old rd1; **Figure 4c**). Like in control animals increase in changepoint rate was dependent on flash intensity (**Figure 4d**, p = 0.021 and 0.004, df = 2, χ^2^= 7.70 and 11.047, n = 56 and 84 trials, kruskalwallis test respectively “young” and “old” groups).

We next used the looming stimuli in order to assess visual acuity. We focused on a reduced spatial frequency range (0.017, 0.068, 0.272 cycles/degree). Both age groups failed to exhibit significant responses to any of the stimuli presented (**Figure 4e,f**; “young”: p = 0.689, 0.894, 0.141, n = 56 trials; “old”: p = 1, 0.445, 0.586, n = 84 trials; signtest at 0.017, 0.068, 0.272 cycles/degree). These negative results are consistent with data from both visual water task and optomotor assays showing loss of spatial discrimination in this genotype (Abdejalil et al., 2005; Thyagarajan et al., 2010).

We wondered whether behavioural responses to spatially structured stimuli such as our looming gratings could be observed in animals with less complete retinal degeneration. To investigate this possibility we used rd12 animals, which carry a mutation in the Rpe65 gene that disrupts recycling of the chromophore (Pang et al., 2005). Previous results based on optokinetic reflex revealed detectable responses below 0.2 cycles/degrees in adult animals aged between 9 and 18 weeks (Wright et al., 2014).

We presented looming grating stimuli over a range of spatial frequencies (0.017-0.272 cpd) to an age-matched cohort (n = 7; 14.4±4.4sd weeks) of rd12 mice. Consistently with published data using the optokinetic reflex, we also found, below the reported limit of 0.2 cycles/degree, significant responses at 0.068 cycles/degrees (**movie 7**, **Figure 4g**, **left panel**; p = 0.009, n = 98 trials; signtest) and a similar response at lowest frequency tested that however did not reach significance (**Supplementary Figure 1a**; p = 0.56, signtest, n = 98 trials, signtest; spatial frequency = 0.017 cycles/degree). Additionally we also found significant responses at 0.272 cycles/degree (**movie8**, **Figure 4g**, **right panel**; p = 0.004, n = 49 trials, signtest). Responses to all stimuli consisted of a reduction in activity similar to the responses observed in control animals. Changepoint results were consistent across stimuli with a larger fraction of trials exhibiting an increase in changepoint rate (51%, 57%, 53% respectively at 0.017, 0.068, 0.272 cycles/degree) compared with trials exhibiting a decrease (34%, 34%, 36%). However reproducibility was lower than in visually intact animals (p = 0.008, permutation test comparing distributions of Δ#chp/s for visually intact and rd12 animals across all spatial frequencies) and the average increments in changepoints were only significant at 0.068 cycles/degree (**Figure 4h**).

### Behavioural Habituation in Mice with Intact and Degenerate Retinas

As animals are presented with multiple repeats of the same or similar stimuli with no associated reward or punishment the salience of those stimuli is expected to “wash off”, reducing the drive for behavioural responses (Rankin et al., 2009). Indeed habituation to looming stimulation has been previously reported (Shang et al., 2018). As habituation could potentially limit the number of effective trials that can be presented to an individual animal we asked to what extent this phenomenon affected our tests. To quantify habituation we focussed on those stimuli and animal groups associated with clearly detectable behavioural responses as with little (e.g. flash stimuli in “old” rd1, **Figure 4**) or no responses this effect is not measurable (e.g. looming stimuli in rd1, **Figure 4e,f**). Note that in order to pre-emptively mitigate the possibility of habituation by reducing expectation the presentation order of the stimuli throughout our study was block randomised (See Methods).

During flash experiments changes in speed in animals with intact vision and in young rd1 were both trending towards lower amplitude in “late” trials (6^th^ and 7^th^ repetitions of the stimulus) compared with those recorded in “early” trials (1^st^ and 2^nd^ repetitions; **Figure 5a**; p = 0.049, 0.039, respectively visually intact and young rd1, permutation test). During looming experiment results from visually intact and rd12 animals were more clearly different. The former did not exhibit significant habituation (**Figure 5c, left panels**; p = 0.482, permutation test) while the latter expressed a substantial reduction in speed changes across trials (**Figure 5c, right panel**; p = 0.001). None of these effects were observed in changepoint rates indicating that while the amplitude of behavioural responses is reduced in “late” trials those responses are still present (**Figure 5b,d**; visually intact: p = 0.718, 0.556 respectively flash and looming; rd1: p = 0.283; rd12: p = 0.401; permutation tests). Overall our data indicate that habituation can have measurable short-term effects over the course of an experimental session.

**Figure 5:**
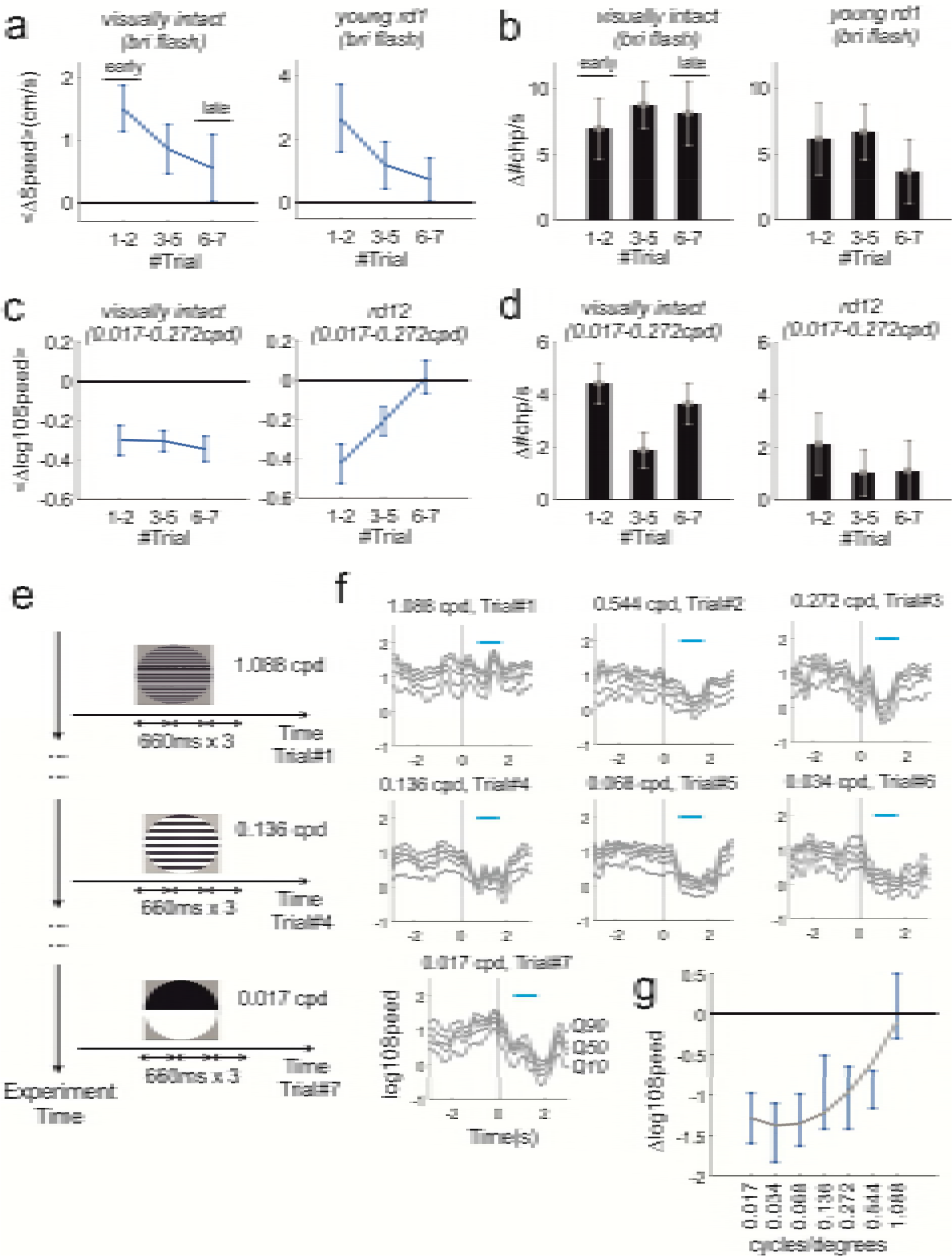
Behavioural habituation to repeated stimulation in visually intact and retinally degenerate mice. **a)** Average changes in speed after stimulus onset as function of trial order for the highest intensity flash (data shown as mean±sem). Represented in left and right panels are visually intact and “young” rd1 animals. Both groups exhibit a negative trend (visually intact: p = 0.049, n = 36 trials; young rd1: p = 0.039; n = 24 trials; comparisons between early and late trials, permutation test). **b)** Difference between changepoint rate after and before stimulus onset (Δ#chp/s) as function of trial order during highest intensity flash for visually intact and young rd1 animals (respectively left and right panel). Neither group showed a significant trend (visually intact: p = 0.718, n = 36 trials; young rd1: p = 0.283; n = 24 trials; comparisons between early and late trials, ranksum test). **c,d)** Like panels **a** and **b** but for looming responses in visually intact and rd12 animals. No significant trend was observed for visually intact animals (panel c: p = 0.482, n = 39 trials; panel d: p = 0.556, n = 39 trials). The rd12 groups exhibited gradual reduction in speed (p = 0.001, n = 21 trials) while no clear trend could be observed in changepoints (p = 0.401, n = 21 trials). **e)** Alternative protocol based on systematic reduction of spatial frequency from highest to lowest (1.088, 0.544, 0.272, 0.136, 0.068, 0.034, 0.017 cycles/degree). Each spatial frequency was presented only on one trial and repeated three times during that trial as in (Yilmaz and Meister, 2013). **f)** Average speed response for all spatial frequencies tested (n = 5 animals, 1 trial/animal). **g)** Polynomial fit for changes in speed as function of spatial frequency (polynomial degree = 2; R^2^= 0.263; same dataset shown in from panel **f**, data shown as mean±sem). Consistent with results obtained by using the block randomised protocol (**Figure 3a-c**) reduction in speed can be observed up to 0.544 cycles/degrees but not at 1.088 cycles/degrees.

In situations where many stimuli have to be presented, an alternative solution to counteract habituation could be to present stimuli in order by starting with those least likely to be detected. To test this possibility we repeated the estimation of spatial acuity in a new cohort of visually intact animals. We selected a wider set of spatial frequencies and presented them in decreasing order (**Figure 5e**). In this way the minimally detectable stimulus (in this case threshold spatial frequency) will be associated with the strongest, non-habituated response, providing the clearest upper bound to estimate visual function. Consistently with the results in **Figure 3a,b** this alternative protocol returned a limiting spatial acuity between 0.5 and 1 cycles/degrees (**Figure 5f,g**).

## Discussion

Our results provide a proof of principle that systematic assessment of visually guided innate responses, such as freeze or escape, represents a promising tool to asses visual capabilities both in mice with intact visual system and in mice affected by different levels of visual impairment.

In the last few years, interest in visually driven innate behaviours has come of age (see e.g. (Evans et al., 2018; Shang et al., 2015a; Wei et al., 2015a)). The growth in interest in such behaviours has been attributable primarily to their application in studies of sensorimotor transformations and action selection. As these behaviours rely upon detection of a visual stimulus, and are readily elicited in untrained animals, they also have clear potential as a basis for characterising visual capacity. However several potential barriers stood in the way of realising that potential at the start of this project: 1) the degree to which behaviourally salient stimuli used to evoke innate responses could be parameterised was unknown; 2) reliable, automated and objective methods for measuring innate behavioural responses were not well defined; 3) the intrinsic variability of innate behaviours may make them too unreliable as an indicator of whether a mouse had detected a visual stimulus; 4) innate responses could quickly habituate; 5) the extent to which visually driven behaviours appear in animals with visual impairment was unexplored. Below, by using our methodological and behavioural results as well as other recent studies, we discuss these issues:

### 1) Parametrising behaviourally salient visual stimuli

Most studies on visually driven innate responses have focussed on looming stimuli in the form of a dark, enlarging, spot. While variants of this stimulus have been considered (see e.g. (Yilmaz and Meister, 2013)) the general conclusion that defensive responses are strongly selective might have prevented a full exploration of the space of effective stimuli. Very recently it was shown that escape response to dark looming exhibits a smooth dependence on the contrast of a looming spot, indicating that negative contrast can be effectively parametrised (Evans et al., 2018). Here we show that darkening looming stimuli do not represent the only stimulus class able to trigger defensive responses: isoluminant stimuli – black and white gratings whose average luminance matches a grey background - can reliably evoke freezing (**Figure 1d**, **Figure 3**). Importantly we have shown that altering the spatial frequency of these gratings can be used to describe mouse spatial acuity. Moreover it is now reasonable to assume that changes in other grating characteristics e.g. contrast, colour etc. could be applied to quantify other visual features.

Although mice with advanced degeneration did not respond to gratings (rd1, **Figure 4a-f**), we found that they did respond to another parameterisable stimulus: full field flashes. Responses to full field flashes have been previously shown to evoke behavioural responses whose magnitude depends on flash intensity (Liang et al., 2015). Unexpectedly the responses we observed, consisting in sharp transient increases in activity, were different and opposite from the behavioural arrests observed by Liang and co-workers (Liang et al., 2015). We note however that the environment in which those behavioural arrests were recorded, a narrow corridor, was substantially different from the open field arena used for this study. Therefore, a likely explanation for different behavioural outcomes is that different environments trigger diverse behavioural responses. Indeed a similar environmental effect, whereby the presence (or absence) of a shelter triggers escape (or freeze), is well known (see e.g. (Tovote et al., 2016)).

### 2) Methods for measuring innate responses

In this work we introduce a new simple method to track multiple body points and establish changepoint analyses to detect innate responses in behavioural time series. These methods allow detection of large changes in behaviours such as sudden locomotion arrests or initiations (see e.g. **Movie 2&7**) as well as more subtle behavioural responses (see e.g. **Movie 1 & 4**). Changepoint analysis is based on PELT algorithm and provides the optimal segmentation under a simple cost function (Killick et al., 2012). It has only one free parameter, which can be quickly estimated from the data with the CROPS algorithm (Haynes et al., 2017) as shown in Methods, therefore results can be easily reproduced. Changepoint analysis is particularly useful for measuring how reliably behavioural responses occur across trials by simply subtracting the rate of changepoint before and after stimulus onset.

### 3) Intrinsic variability of innate behaviours

Spontaneous behaviours can vary substantially between individuals. Thus, different animals exposed to the same threating visual stimulus can employ diverse, and sometimes opposing (e.g. freezing or escape), behavioural strategies. However recent results indicate that different classes of visual stimuli can bias towards different types of innate responses and this bias is shared across individuals. Thus De Franceschi (De Franceschi et al., 2016) has shown that looming and sweeping dark spots can consistently drive mice to select either escape (for a looming spot) or freezing (for a sweeping spot). In this paper we describe another clear behavioural dichotomy as the same group of animals express sharp increments in speed after a flash presentation and freezing-like behaviours under loom+gratings stimuli (respectively **Figure 2a,3a**). Moreover changepoint analysis can automatically detect for both negative and positive changes in speed and therefore also addresses intrinsic variability of innate responses.

### 4) Innate responses could quickly habituate

One obvious potential solution to inherent behavioural variability is to record responses to multiple stimulus presentations. However, when animals are presented with repeated sets of threatening stimuli, that turn out to be inconsequential, behavioural responses will quickly habituate (Rankin et al., 2009). Therefore habituation could represent a barrier to presenting large numbers of similar stimuli to the same animal as a method of addressing response variability and in order to describe responses to quantitative alterations in the stimulus (e.g. when estimating threshold spatial acuity). While we do observe some level of habituation in the amplitude of behavioural responses (**Figure 5a,c**) those were still present across multiple repetitions (up to 7) and detectable by changepoint analysis (**Figure 5b,d**). Thus when multiple stimuli must be presented to estimate detection thresholds, we find that habituation can be addressed either by randomising the presentation order of a limited set of stimuli (**Figure 3**) or by presenting a wider set that starts from the least detectable stimulus (**Figure 5e-g**).

In this study we did not investigate the potential for long-term habituation to the same set of stimuli presented on different experimental sessions. Others have described this phenomenon and reported it to be stimulus specific (Rankin et al., 2009). Our data are consistent with this view as we were able to record responses to two different sets of stimuli (flash and looming gratings) from the same visually intact animals on different days (**Figures 2&3**). Thus, while it may not be sensible to re-test the same stimuli on a different occasion, simple modifications of the stimuli (e.g. changing orientation of looming gratings) that have been shown to enhance perception of novelty (Cooke et al., 2015) could allow re-testing.

### 5) To what extent visually driven behaviours are conserved in animals with different levels of visual impairment?

A fundamental concern about the use of visually driven innate responses in describing sight loss is the extent to which they are conserved in animals with impaired visual function. Clearly an all-or-none situation whereby innate behaviours are either reliably expressed or fully abolished would only allow for a binary discrimination between animals with intact or impaired function. Given the wide spectrum of visual impairments occurring in different mouse models of ocular diseases and individual variability in degeneration and treatment efficacy this kind of binary discrimination would be of little use. In this study we show that visually driven innate responses allow for a finer discrimination across different levels of visual impairments. Thus, quantitative differences (in flash responses between rd1 age groups, **Figure 4a**) as well as qualitative differences (presence or absence of freeze responses to spatial stimuli, **Figure 4e-h**) allowed for discrimination between severity and type of retinal degeneration.

## Conclusions

Our results indicate that systematic assessment of innate responses elicited by parameterisable stimuli represent a viable approach for measuring visual capabilities both in mice with intact visual function and in mouse models of retinal degeneration. Furthermore innate behaviours have several advantages over other established tests of visual function. The tests we propose are quick; can be performed on naïve mice; do not require specialist training for the experimenter; and the analyses are all automated. Compared with tests based on brainstem driven reflexes, such as optokinetic nystagmus or optomotor following, they capture higher limits in spatial acuity that match those obtained with the visual water task. However differently from the visual water task they do not require preliminary training sessions that can last several weeks in animals with limited vision (Thyagarajan et al., 2010), can cause significant distress (Vorhees and Williams, 2014) and introduce further variability in the results due to the learning process itself (Coppens et al., 2010; Flagel et al., 2009). Finally, the use of innate behaviours allows for recording a wider repertoire of behavioural responses, such as speed increments after full field flashes and freeze after looming, increasing the experimenter’s ability to discriminate different levels of visual function.

## Supplementary Information

**Supplementary Figure 1:**
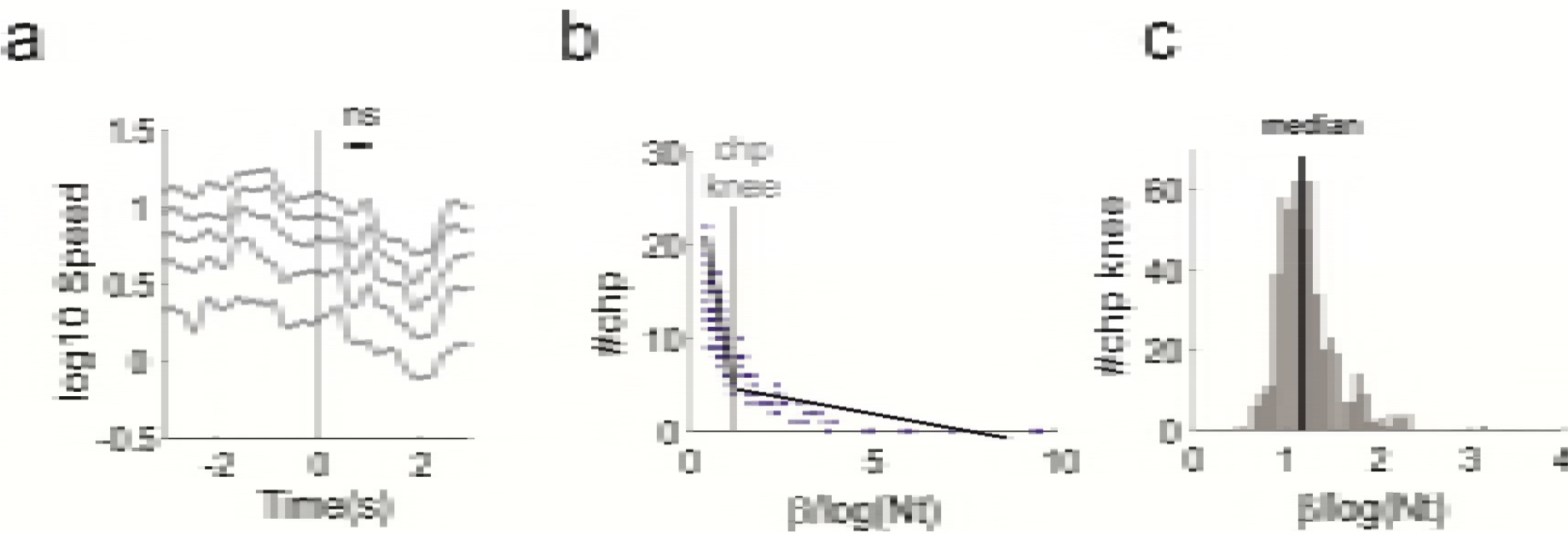
Looming response at 0.017 cpd for rd12 animals & data driven estimation of penalty value for changepoint detection. **a)** Movement response, in log scale, to 0.017 cycles/degree gratings. **b)** Individual trial example where the number of changepoints (“#chp”, blue dots) across the dimensions of the multivariate time series are plotted as function of penalty. The knee (“chp knee”, grey vertical line) is estimated by fitting a segmented function (black lines meeting at the knee point, see Methods) between number of changepoints and penalty values. **c)** Distribution of knee values for our dataset. The median (black line) is then used for all further changepoint analyses.

## Methods

### Ethical Statement

Experiments were conducted in accordance with the Animals, Scientific Procedures Act of 1986 (United Kingdom) and approved by the University of Manchester ethical review committee.

### Experimental Set-Up

We used an open field arena and recorded the animals with 2 programmable global shutter cameras (Chamaleon 3 from Point Grey) stably mounted on optical rails (Thorlabs; **Figure 1a**). In order to avoid saturation due to changing light levels in the visual stimuli the camera lenses were covered with infrared cut-on filters (Edmund Optics) and fed with constant infrared light. Visual stimuli were delivered by a projector onto a rear projection screen. The flashes for the full field contrast stimuli were provided by two LEDs mounted inside the arena (LED engin LZ4-00B208; controlled by T-Cube drivers, Thorlabs). The experiments were controlled by using Psychopy (Peirce, 2007). Frame acquisition was synchronized with the projected images and across cameras by a common electrical trigger delivered by an Arduino Uno board (arduino.cc). All stimuli were calibrated by using a spectroradiometer (Bentham Instruments) and retinal irradiance values for each photoreceptor were calculated by using Govardovskii templates (Govardovskii et al., 2000) and lens correction functions (Jacobs and Williams, 2007) as previously described (Storchi et al., 2017; Storchi et al., 2015). Summary information for all stimuli are provided in **Table 1&2**.

For all experiments we delivered 7 blocks of 2-3 different stimuli whose order was independently randomized within each block. We set inter-stimulus interval at 73 seconds. For each trial the video recording started 6 seconds before and ended 7 seconds after the stimulus onset. Framerate was set at 15Hz and allowed us to collect 200 frames per trials.

### Animal Housing & Handling

All mice were stored in cages of 2-4 individuals and were provided with food and water ad libitum. During transfer between the cage and the behavioural arena we used the tube handling procedure instead of tail picking, as prescribed in (Hurst and West, 2010), in order to minimise stress and reduce variability across animals.

#### Immunohistochemistry & cell imaging

Mice were culled, enucleated and eyes pierced with 24gauge needle before being transferred to 4% PFA in PBS and stored overnight at 4C°. Retinas were then dissected and permeabilised in PBS with 1% Triton-X for 3 × 10mins. A background block incubation in PBS with 1% TritonX with 10% Donkey serum was conducted before retinas were incubated in primary antibody solution (PBS with 1% Triton-X with 2.5% donkey serum with 1:250 dilution of rabbit anti-SWS cone opsin antibody, AB5407 Merck Millipore) overnight at 4°C. Retinas were then washed in PBS with 0.2% TritonX for 4 × 1 hour, then incubated in secondary antibody solution (PBS with 1% TritonX and 2.5% Donkey serum with 1:250 dilution of donkey anti-rabbit Alexa 546, A10040 Thermo Fisher) overnight at 4°C. Retinas were washed in PBS with 0.2% TritonX for 4 × 1 hour, then washed in distilled water for 10mins before being mounted ganglion cell-side up on microscope slides using Prolong Gold anti-fade mounting reagent (Thermo Fisher) and left to cure overnight at room temperature.

Images were collected on a Leica TCS SP5 AOBS inverted confocal using a 60x objective. The confocal settings were as follows, pinhole [1 airy unit], scan speed [1000Hz unidirectional], format [1024 × 1024]. Images were collected using PMT detectors with the following detection mirror settings; Texas red 602-665nm using the 594nm (100%) laser line. When acquiring 3D optical stacks the confocal software was used to determine the optimal number of Z sections. Only the maximum intensity projections of these 3D stacks are shown in the results.

### Mouse Tracking

We first isolated the mouse body by using a static background acquired at the beginning of each experiment before introducing the animal into the arena. The initial subtraction was refined by an opening operation (erosion followed by dilation) that removed mouse excrements that occasionally accumulated on arena floor during the experiment. These operations allowed us to extract a tightly bounded box around the animal body that sped up subsequent calculations.

We found that the classic Harris corner detector was effective in detecting most relevant landmarks such as ears, snout tip, paws extremities, tail base and whiskers. However when just applied to the original image it only returned a partial list of features per image. In order to increase the yield of Harris detector we applied it to the associated 6 levels image pyramid representation (sample factor = 1.25 with 13 points Gaussian smoothing, σ = 1.25). This returned a dense labelling of body landmarks as shown in **Figure 1b** (bottom panels).

In order to quantify landmark speed the landmarks between two successive images were initially matched by using Pearson’s correlation. Typically not all matches were correctly assigned and incorrect assignments could introduce artefactual jumps in the behavioural time series. To address this problem we applied the following algorithm to identify and reject outliers. We estimated a partial Procrustes superimposition (rotation + translation) between matched landmarks coordinates. The estimation was performed by employing the RANSAC algorithm (Fischler and Bolles, 1981) that allowed us to reject outliers given a fixed error threshold (inlier threshold = 10 pixels). While a single linear transformation was effective in capturing movements with consistent direction and amplitude across the whole mouse body (**Figure 1b**, right panel) it failed to capture movements where different body parts where displaced in different directions and/or with different amplitudes. In order to capture the whole gamut of mouse movements we extended our approach by combining clustering and Procrustes analysis. Thus landmark coordinates were clustered via the *kmeans++* algorithm (Arthur and Vassilvitskii, 2007) by systematically varying K, the number of clusters (K = [1,5]; for each value of K clustering was performed 10 times to avoid suboptimal solutions). Procrustes partial superimposition was then separately estimated for each cluster (**Figure 1b**, centre panel). To identify the optimal number of clusters we then compared the residuals by using as criterion the Minimum Description Length (Rissanen, 1978) that allowed us to incorporate model complexity (i.e. the number of clusters) and the cost of different number of outliers. The description length of each model was defined as:

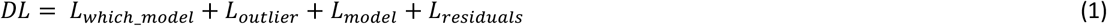

and

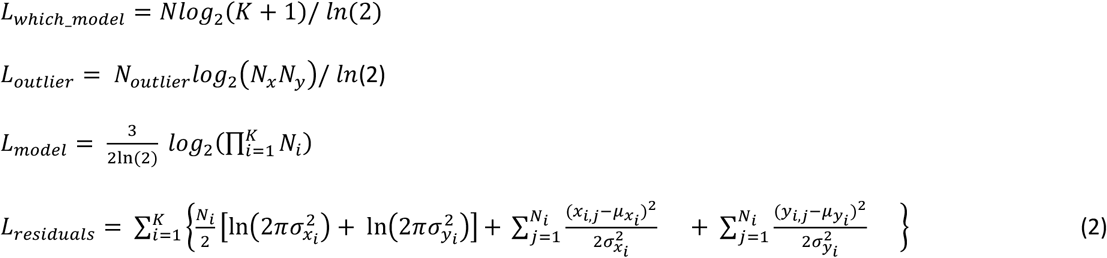

where *N*_*x*_*N*_*y*_ represents the area of the bounding box around the animal, *N*_*outlier*_ the total number of outliers, *N*_*i*_ the number of inliers for the *i*^*th*^ cluster, *x*_*i*,*j*_ the *j*^*th*^ residual of the *i*^*th*^ cluster on the x direction. The first term of equation 1, *L*_*which_model*_ accounts for the number of *nats* (natural bits) required to identify each point as outlier or belonging to the *i*^*th*^ cluster. The second term costs the outliers as bounded integers in the area bounding box. The third term penalizes for the number of clusters and the factor 3 corresponds to the number of parameters required to define the partial Procrustes superimposition for each cluster (1 for rotation, 2 for translation). The final term uses the fact that the cost of encoding residuals can be approximated by the negative log likelihood.

### Generation of Movement Time Series

We used the distribution of landmarks speed to generate multidimensional behavioural time series. We first calculated the speed of each landmark between two consecutive frames (**Figure 1b**, bottom panels). From the distribution of landmarks speed we then calculated the 10^th^, 30^th^, 50^th^, 70^th^, 90^th^speed quantiles for each frame (**Figure 1b**, top panels; respectively Q10 to Q90). Each of these quantiles, estimated separately for each of the two cameras used (**Figure 1a**), defined a dimension of the multivariate time series (**Figure 1c**). For all subsequent analyses the quantiles were then averaged across the cameras (**Figure 2a, 3a, 4a,e,g**).

### Changepoints Detection

In order to find the number and the location of the changepoints we used the Pruned Exact Linear Time method (PELT) (Killick et al., 2012). PELT is designed to minimize a penalised cost function of the following form:

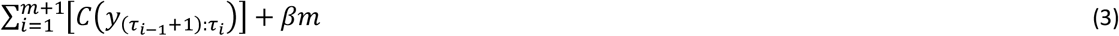

Where *C* is the cost function, *τ* represents the changepoints, *m* is the number of changepoints and β is the penalty for each additional change. This method has two main advantages: it calculates the global optimum of (3) and it does so in a computational cost, under mild conditions, that scales linearly with the number of data points. We define the cost function *C* as follows:

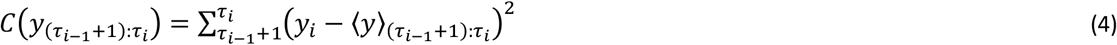

Where <y> represents the average speed between two successive changepoints. The β term in (3) represents a penalty against overfitting. Popular choices for the penalty function include the Schwarz Information Criterion (Schwarz, 1978). Several authors also proposed different solutions to this problem by adaptations of the SIC to the problem of multiple changepoint analyses (Zhang and Siegmund, 2007). As a general consensus lacks about which criterion is more convenient for a given dataset we devised a method to automatically derive a suitable penalty range from the data. Therefore, instead of specifying a penalty value, for each trial we systematically scanned across a wide range of β values by using the Changepoint for a Range Of PenaltieS (CROPS) algorithm (Haynes et al., 2017). The CROPS algorithm efficiently finds all the sets of changepoints whose segmentations are optimal under some choice of β within an interval [βmin, βmax] by sequentially dividing this interval until guaranteed convergence. We then used the output of this algorithm, the number of changepoints as function of penalty values, to fit a piecewise linear model represented by two lines intersecting at a “knee” point (**Supplementary Figure 1b**). The knee point marks the transition between under and over fitting and thus represents an ideal choice for the penalty. The segmented relationship between the number of changepoints and the penalty value β was modelled following (Muggeo, 2003) as

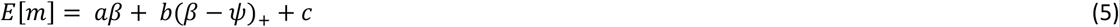

where parameters *a* and *b* represent respectively the slope of the left line segment and the difference in slopes between the left and the right line, (*β* − *ψ*)_+_equals 1 for *β* > *ψ* and 0 otherwise, and *ψ* represents the knee point. By using the iterative estimation proposed by (Muggeo, 2003) we then obtained a maximum likelihood estimation of all the parameters including the knee point (**Supplementary Figure 1b**). We repeated this procedure across all trials to obtain the distribution of knee values for our datasets that provided a suitable range for choosing the penalty value β (**Supplementary Figure 1c**).

#### Statistics

Statistics derived from tracking data were applied either to the raw multivariate time series (for full field flash) or to a truncated log transformation of these data (for loom+gratings). The former was positively skewed, the second negatively skewed. Accordingly changepoints applied to the raw data was more biased towards detecting positive changes in movement, such as flights, the second towards negative changes such as freezing.

The significance of behavioural responses to individual stimuli was assessed by using the signtest (in **Figure 2a, 3a, 4a,e,g**). The test statistic was obtained by subtracting the mean landmarks speed after stimulus onset from that recorded before the stimulus onset. Speed values were estimated in time windows of equal duration (0.53 seconds for both looming and flash). Comparison between flash responses was performed on the same speed distributions by using the ranksum test (e.g. between “young” and “old” rd1 mice, **Figure 4a**). Statistical significance for changepoints was calculated by subtracting the rate of change points before - *#chp/s(pre)* - and after - *#chp/s(post)* - stimulus onset using the same time windows and applying a ranksum test. Thus the difference in changepoint rate (Δ#chp/s in **Figure 2b, 3b, 4d,f,h**) was calculated as Δ#chp/s = *#chp/s(pre) - #chp/s(post).* Statistical significance of intensity dependence for flash responses shown in **Figure 2b, 4d** was calculated from the same data by using kruskalwallis test.

In order to test habituation both in speed and changepoint responses we used permutation tests. First the difference in speed responses (or changepoint responses) was calculated between early and late trials (**Figure 5a,b**). Then this difference was recalculated 100000 times by shuffling the order of the stimulus trials and a p-value was determined as the fraction of instances in which these surrogate values were larger than the original statistic.

### Software Resources

Changepoint analyses were performed in R (2014). The code for changepoint analysis, including the CROPS method, can be found in the ‘changepoint’ package (Killick and Eckley, 2014). For knee estimation we used the ‘segmented’ package (Muggeo, 2008). The code for mouse tracking was written in MATLAB (Matworks, Natick, Massachusetts, USA).

## Acknowledgements

We thank Annette Allen for thorough discussions during the preparation of the manuscript; Sumayyah Zanna Muhammad for helpful feedback during the early phase of the project; and Robin Ali at University College London for supplying us with the rd12 mice. The work was funded by National Centre for Replacement Refinement and Reduction of Animals in Research (NC3Rs) via a David Sainsbury Fellowship (NC/P001505/1) to R.S. and a project grant from the Medical Research Council to R.J.L. (MR/N012992/1).

## Movie Captions

**Movie 1:** Increase in movement by parts of the mouse body; here occurring as a change in rearing posture elicited by “bright” flash (stimulus occurring at time 0). Top left panel represents the quantile of the speed distribution (10^th^, 30^th^, 50^th^, 70^th^, 90^th^ quantiles - respectively Q10 to Q90) for the two cameras used for the study (for ease of representation here only the movie from camera1 is shown). Bottom left panels represents single changepoints associated with quantile time series (Q10 to Q90 for the two cameras). Pooled changepoints (see Methods) are also reported at the top line (“Pooled”).

**Movie 2:** Increase in full body movements elicited by “bright” flash; here the flash initiates locomotor activity. Panels displayed as in Movie 1.

**Movie 3:** Decrease in full body movements elicited by loom+gratings stimulus (gratings at 0.272 cycles/degree); here the gratings (occurring at time 0) arrests mouse locomotion in freeze-like behaviour. Panels displayed as in Movie 1.

**Movie 4:** Decrease in movements by parts of the mouse boy elicited by loom+gratings stimulus (gratings at 0.0.068 cycles/degree); here the animal, engaged in stationary exploration, produces a freeze-like response. Panels displayed as in Movie 1.

**Movie 5:** Increase in movement in a “young” rd1 mouse elicited by “bright” flash. Panels displayed as in Movie 1.

**Movie 6:** Increase in movement in an “old” rd1 mouse elicited by “bright” flash. Panels displayed as in Movie 1.

**Movie 7:** Decrease in movement in an rd12 mouse elicited by loom+gratings stimulus (gratings at 0.068 cycles/degree). Panels displayed as in Movie 1.

**Movie 8:** Decrease in movement in an rd12 mouse elicited by loom+gratings stimulus (gratings at 0.272 cycles/degree). Panels displayed as in Movie 1.

